# Multifocal Ectopic Purkinje Premature Contractions due to neutralization of an *SCN5A* negative charge: structural insights into the gating pore hypothesis

**DOI:** 10.1101/2024.02.13.580021

**Authors:** Andrew M. Glazer, Tao Yang, Bian Li, Dana Page, Mohamed Fouda, Yuko Wada, Megan C. Lancaster, Matthew J. O’Neill, Ayesha Muhammad, Xiaozhi Gao, Michael J. Ackerman, Shubhayan Sanatani, Peter C. Ruben, Dan M. Roden

## Abstract

**Background:** We identified a novel *SCN5A* variant, E171Q, in a neonate with very frequent ectopy and reduced ejection fraction which normalized after arrhythmia suppression by flecainide. This clinical picture is consistent with multifocal ectopic Purkinje-related premature contractions (MEPPC). Most previous reports of MEPPC have implicated *SCN5A* variants such as R222Q that neutralize positive charges in the S4 voltage sensor helix of the channel protein Na_V_1.5 and generate a gating pore current.

**Methods and Results:** E171 is a highly conserved negatively-charged residue located in the S2 transmembrane helix of Na_V_1.5 domain I. E171 is a key component of the Gating Charge Transfer Center, a region thought to be critical for normal movement of the S4 voltage sensor helix. We used heterologous expression, CRISPR-edited induced pluripotent stem cell-derived cardiomyocytes (iPSC-CMs), and molecular dynamics simulations to demonstrate that E171Q generates a gating pore current, which was suppressed by a low concentration of flecainide (IC50 = 0.71±0.07 µM). R222Q shifts voltage dependence of activation and inactivation in a negative direction but we observed positive shifts with E171Q. E171Q iPSC-CMs demonstrated abnormal spontaneous activity and prolonged action potentials. Molecular dynamics simulations revealed that both R222Q and E171Q proteins generate a water-filled permeation pathway that underlies generation of the gating pore current.

**Conclusion:** Previously identified MEPPC-associated variants that create gating pore currents are located in positively-charged residues in the S4 voltage sensor and generate negative shifts in the voltage dependence of activation and inactivation. We demonstrate that neutralizing a negatively charged S2 helix residue in the Gating Charge Transfer Center generates positive shifts but also create a gating pore pathway. These findings implicate the gating pore pathway as the primary functional and structural determinant of MEPPC and widen the spectrum of variants that are associated with gating pore-related disease in voltage-gated ion channels.

## Introduction

Na_V_1.5 (encoded by *SCN5A*) is the major cardiac voltage-gated sodium channel. Variants in *SCN5A* are associated with multiple arrhythmia disorders, including loss of function diseases (Brugada syndrome and progressive cardiac conduction defect) and long QT syndrome (enhanced late current).^1^ *SCN5A* mutations have also been associated with the rarer phenotype of multifocal ectopic Purkinje-related premature contractions (MEPPC).^2-4^ MEPPC presents with frequent junctional or ventricular ectopic beats, often early in childhood, and frequently leading to dilated cardiomyopathy. MEPPC is important to recognize since suppression of the frequent ectopy can reverse the associated cardiomyopathy.^2-4^

Voltage gated sodium channels including Na_V_1.5 pass through the plasma membrane 24 times, forming 4 major domains (I through IV). Within each domain, transmembrane helices S1-S4 contribute to voltage sensing and S5 and S6 largely form the central pore through which sodium passes. The S4 helices contain positively charged residues (arginines or occasionally lysines) at every third residue. The S4 helix moves in response to changes in membrane voltage and this movement underlies the channel’s voltage sensitivity. Charge neutralizing mutations of the voltage-sensing positively charged residues (arginines) of the S4 helices are the most common *SCN5A* variants associated with MEPPC.^5^ The most common variant in this class is R222Q,^2-4^ and R219H,^6^ R225W, R225P,^7^ and R814W^8^ have also been reported. These mutations are thought to generate a second (non-physiologic) ion permeation pathway in the S1-S4 gating region, leading to a “gating pore” current or omega current.^6,9,10^ Several additional variants that are not in S4 arginines have been described in MEPPC (A204E,^11^ L828F,^12^ G213D,^13^ and M1851V^14^) and these variants are mainly thought to act through alternate mechanisms such as shifts in the voltage-dependence of channel activation and/or inactivation generating an increased “window current”.

Studies in the Drosophila *Shaker* potassium channel identified a “gating charge transfer center” (GCTC), critical for movement of the voltage sensor; key features included positively charged residues in S4, two negatively charged residues in S2 and S3, and a key phenylalanine (tyrosine in the analogous position in Na_V_1.5).^15,16^ This motif is highly conserved across voltage-gated ion channels, and mutations at S4 arginines have been reported not only in MEPPC but also in sodium and calcium channel-related hypokalemic and normokalemic periodic paralysis, familial hemiplegic migraine, malignant hyperthermia, and familial epilepsy.^15^ It is thought that the GCTC helps form a barrier that separates intracellular and extracellular water-filled crevices. Mutations can disrupt this barrier, creating a water-filled channel that generates the gating pore current through a pathway separate from the ion channel’s central permeation pore.^15^

We report a neonate with MEPPC resulting from a novel *SCN5A* variant, E171Q. This variant, unlike other variants that result in a gating pore current, neutralizes a negative charge in the S2 helix of domain I, a previously unrecognized disease mechanism in MEPPC. We use patch clamping to characterize E171Q in heterologous expression and CRISPR-edited iPSC-derived cardiomyocyte models, and identify a large gating pore current generated by the variant channel and blocked by low concentrations of flecainide. We model this variant with molecular dynamics simulations and propose a structural model that explains our findings.

## Methods

The patient’s clinical course is described below (Results).

### Overview of *in vitro* methods

Variants were studied using patch clamping in two *in vitro* systems: (1) Plasmids with *SCN5A* mutations of interest were generated and the mutant cDNAs were expressed in Human Embryonic Kidney 293 (HEK293) cells. (2) Induced pluripotent stem cells (iPSCs) were CRISPR-edited to generate *SCN5A* E171Q and differentiated into cardiomyocytes (iPSC-CMs). iPSCs were obtained from a healthy volunteer under IRB approval as previously described.^17^

### HEK293 cells and plasmid mutagenesis

Electrophysiologic studies were performed independently at laboratories at Vanderbilt University Medical Center (VUMC) and at Simon Fraser University (SFU).

At VUMC, the wild-type plasmid included the SCN5A cDNA, an internal ribosomal entry site, mCherry, and a blasticidin selection cassette (SCN5A wildtype:IRES:mCherry-blasticidinR). The *SCN5A* mutations E171Q (Ensembl transcript ENST00000333535 c.511G>C) and R222Q (c.665G>A) were introduced with a QuikChange Multi Lightning kit (Agilent) with primers 5’ttcaccgccatttacacctttCagtctctggtca (E171Q) or 5’ttacgcaccttccAagtcctccgggcc (R222Q). All cDNA coordinates are presented for Ensembl transcript ENST00000333535, which includes Q1077; however, the wildtype plasmid used in this study has a deletion of Q1077 to match the most common splice isoform in the heart.^18^ We first mutagenized plasmids containing a small region of *SCN5A* (DNA bases 1-727) and verified the mutations by Sanger sequencing. Next, we subcloned the small plasmid with restriction enzymes AgeI and BssHII (New England Biolabs) into a plasmid with full length *SCN5A* and again Sanger sequence verified the plasmids. The resulting plasmids (*SCN5A* wildtype:IRES:mCherry-blasticidinR, *SCN5A* E171Q:IRES:mCherry-blasticidinR, or *SCN5A* R222Q:IRES:mCherry-blasticidinR) were integrated into Human Embryonic Kidney 293 (HEK293) landing pad cells.^19^ Stable lines expressing a high level of the *SCN5A* variant were generated and validated by flow cytometry as we have previously described.^18^

At SFU, the E171Q variant was introduced into an *SCN5A* expression plasmid (pCDNA3.1) using the QuikChange Lightning Site-Directed Mutagenesis kit (Agilent). HEK293T cells were grown at pH 7.4 in filtered sterile DMEM nutrient medium (Life Technologies), supplemented with 5% FBS and maintained in a humidified environment at 37°C with 5% CO2. Cells were transiently co-transfected with the human cDNA encoding the Nav1.5 α-subunit, the β1-subunit, and eGFP. Transfection was done according to the PolyFect transfection protocol (Qiagen). A minimum of 8-hour incubation was required after each set of transfections. The cells were subsequently dissociated with 0.25% trypsin–EDTA (Life Technologies) and plated on sterile coverslips.

### CRISPR editing of iPSC cells

A CRIPSR guide for the *SCN5A* E171Q mutation was designed using the CRISPOR online tool.^20^ We cloned the guide sequence (AATCTTGACCAGAGACTCAA-AGG) into SpCas9-2A-GFP (pX458, Addgene #48138)^21^ by annealing complementary primers 5’CACCGAATCTTGACCAGAGACTCAA and 5’AAACTTGAGTCTCTGGTCAAGATTC, phosphorylating the primers using T4 PNK (NEB) and digesting and ligating the insert into the pX458 plasmid using BbsI (NEB) and T4 Ligase (NEB). This process created a new guide RNA expression plasmid SpCas9-2A-GFP-E171Q. A 150-base single stranded rescue DNA was designed with *SCN5A* genomic sequence with two mutations present: E171Q (c.511G>C) and a synonymous mutation (c.507C>A) to disrupt the PAM site and prevent re-cutting by Cas9. The guide plasmid and the rescue DNA were electroporated into population control iPSC-CMs using a Neon transfection system. After two days, GFP-positive cells were FACS-sorted and plated sparsely. After ∼1 week of growth, single colonies were picked, DNA was isolated using QuickExtract (BioSearch Technologies), and PCR was performed using primers 5’ GTTGAAGGGAGCTTTGTGGG and 5’GAAGCCAGAAAGAGAGGGGT. PCR products were cleaned with ExoSap It (Applied Biosystems) and Sanger sequenced to identify clones with the desired mutations in heterozygous form. iPSC cells were grown and differentiated to iPSC-derived cardiomyocytes as previously described.^17^ Cells were studied 35-40 days post-differentiation. We did not generate R222Q iPSC-CMs because that mutation is in exon 6B, which undergoes alternate splicing; an alternate fetal exon (6A) is the predominant isoform in iPSC-CMs.^22^

### Patch clamping

Detailed electrophysiological methods for voltage- and current-clamping experiments are presented in Supplemental Methods.

### Structural simulation of Na_V_1.5 voltage sensing domain I

We simulated the voltage sensing domain (S1-S4) of domain I (VSD1) for wildtype, E171Q, and R222Q. The R222Q variant was selected as a positive control because it has been shown to result in gating pore current through VSD1 and it is the most common MEPPC mutation.^10^ The wildtype Na_V_1.5 VSD1 was simulated as a negative control as it lacks a gating pore current. For each system, we ran three replications with different initializations of starting atom velocities. Further details of the simulated systems can be found in Table S2.

### Statistical analysis

Data are presented as mean ± standard error. Comparisons between WT and E171Q groups of cells were performed using a t**-**test (two-sample assuming unequal variances), implemented in Excel. Multiple comparisons were performed using Dunnett’s test.

## Results

### Clinical presentation

The proband was born at term to a healthy woman during her first pregnancy. At birth, the proband exhibited salvos of a rapid irregular rhythm (Figure 1A). The native QRS was difficult to define due to a limited number of normally conducted QRS complexes. Most beats were wide complex (for age) with ventriculo-atrial dissociation. There were also episodes of non-conducted sinus beats and rare conducted sinus beats (Figure 1B). The rhythm was predominantly an accelerated wide complex rhythm with runs of ventricular tachycardia of a different morphology (Figure 1C). Occasional periods of Wenckebach atrioventricular conduction were captured. Despite the rhythm, the child remained asymptomatic and thrived, even tolerating an elective craniosynostosis repair. The ejection fraction at birth was 55%, at the low end of the normal range, but by 7 months of age it had fallen to 31%. He was then admitted for a trial of flecainide at 20 mg bid (2 mg/kg/dose). Holter monitoring subsequently demonstrated a reduction in his ventricular ectopy and average rates, and echocardiography showed an improvement in his ejection fraction to 66%. He remains on flecainide and is asymptomatic with an average heart rate on Holter monitor of 100 bpm.

**Figure 1:**
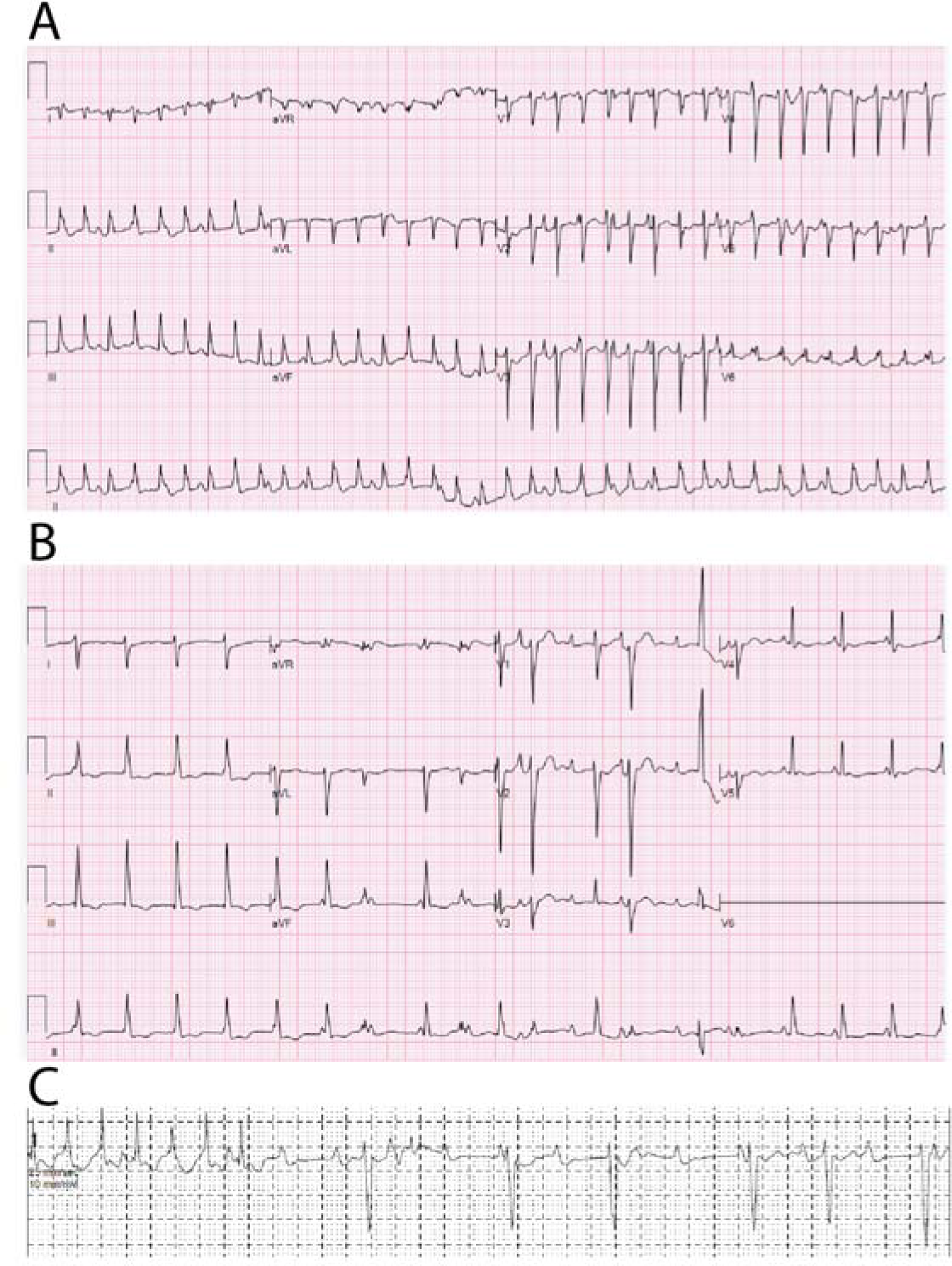
Patient phenotype. A) 12-lead ECG demonstrating AV block dissociation with ventricular tachycardia at ∼214/min. B) 12-lead ECG showing predominant junctional or ventricular ectopy, with a sinus beat (14^th^ QRS). C) Rhythm strip demonstrating both tachycardia as well as bradycardia due to second and third degree AV block.

The proband and parents underwent sequencing using a 217 gene cardiac panel (Blueprint Genetics). Three variants of interest were identified in the proband: (1) *SCN5A* c.511G>C (p.Glu171Gln, referred to here as E171Q), which was *de novo* and classified by the testing company as a variant of uncertain significance (VUS). (2) *FLNC* c.2377G>A (p.Glu793Lys), a paternally inherited VUS. (3) *DMD* c.2911G>A (p.Asp971Asn), a maternally inherited VUS. The parents were phenotypically normal. Based on the phenotypic findings of ventricular tachycardia, cardiomyopathy, and a positive response to flecainide, the patient was diagnosed with multifocal ectopic Purkinje-related premature contractions (MEPPC), suggesting the *SCN5A* variant might be causative. AlphaMissense^23^ predicted that the VUS in *SCN5A* was likely pathogenic while those in *FLNC* and *DMD* were likely benign (Table S1).

### Conservation and computational predictions of *SCN5A* E171Q

E171 is located in Domain I of Nav1.5, a domain which contains most of the known MEPPC-associated variants (Figure 2A). However, E171 is located in the Domain I S2 helix, whereas most of the MEPPC-associated variants alter arginines in the S4 helix. The amino acid homologous to Na_V_1.5 E171 is conserved across all nine voltage-gated sodium channel proteins (Na_V_1.1-1.9), several voltage-gated calcium channels, and the potassium channels *Shaker* and K_V_7.1 (Figure 2B). In the gnomAD v4 dataset of 807,162 individuals, Na_V_1.5 E171Q was not present. In addition, no other missense variants at the E171 position in Na_V_1.5 were observed in gnomAD, in contrast to many other positions in the S2 helix (Figure 2C). The initial study in *Shaker* describing the GCTC identified a key role for a phenylalanine residue at the position analogous to Y168 in Na_V_1.5, another residue with no variants reported in gnomAD (Figure 2C).

**Figure 2:**
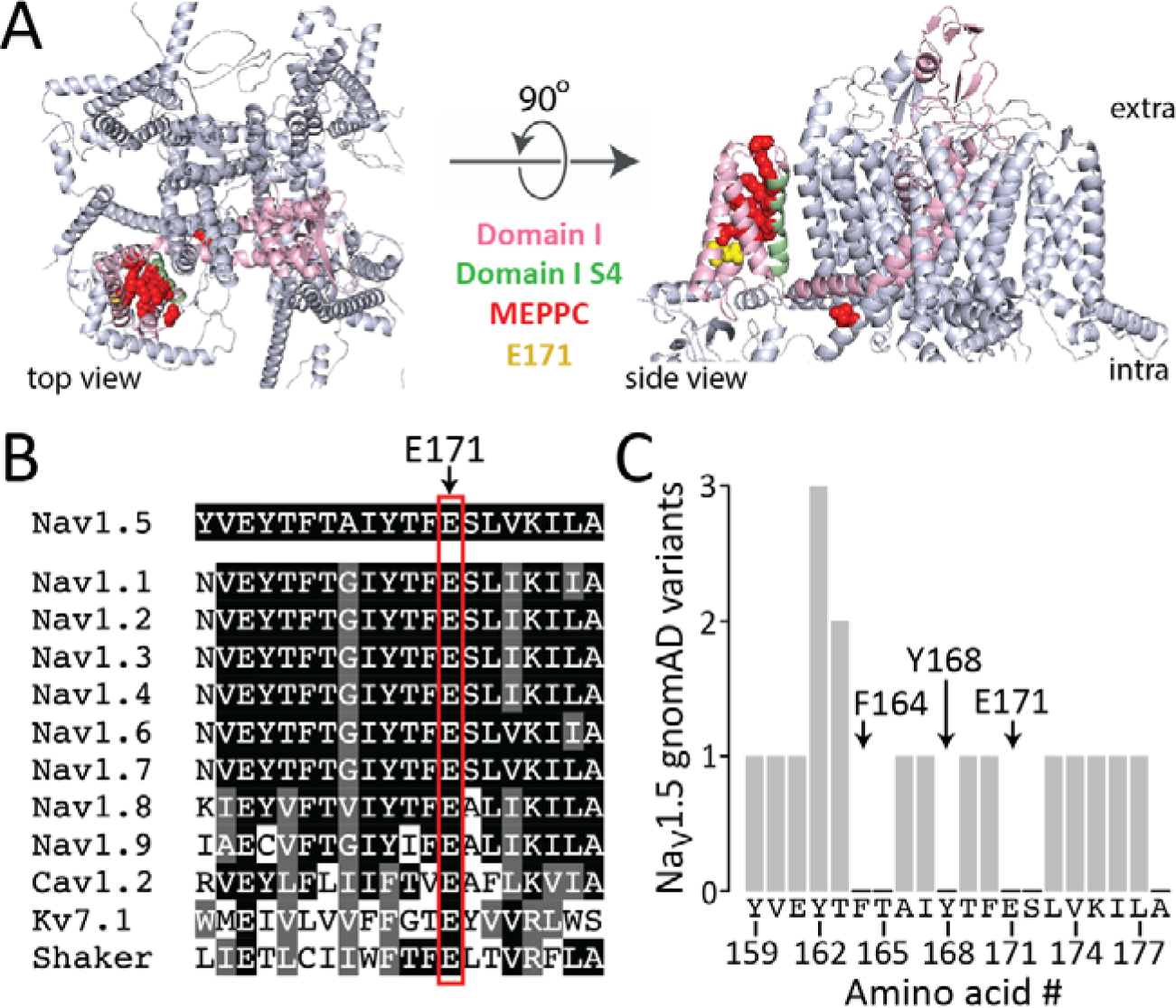
Conservation of Na_V_1.5 E171 and low prevalence of variation in the population. A) Top and side views of Na_V_1.5 structure. Domain I is indicated in pink, the Domain I S4 helix in green, known MEPPC-associated variants in red, and E171 in yellow. B) Multiple sequence alignment of the Domain I, S2 helix of Na_V_1.5 and other voltage gated cation channels. All channels are from humans except for Shaker, which is from *Drosophila melanogaster*. C) Missense variants in the Domain I, S2 helix of Na_V_1.5 in the genome aggregation database (gnomAD v4) in >800,000 individuals. E171 and two hydrophobic residues that also act in the Gating Charge Transfer Center are indicated.

### Na_V_1.5 E171Q generates a late current and a gating pore current in HEK cells

HEK293 cells expressing R222Q or E171Q generated inward sodium current (Figure 3A).

**Figure 3:**
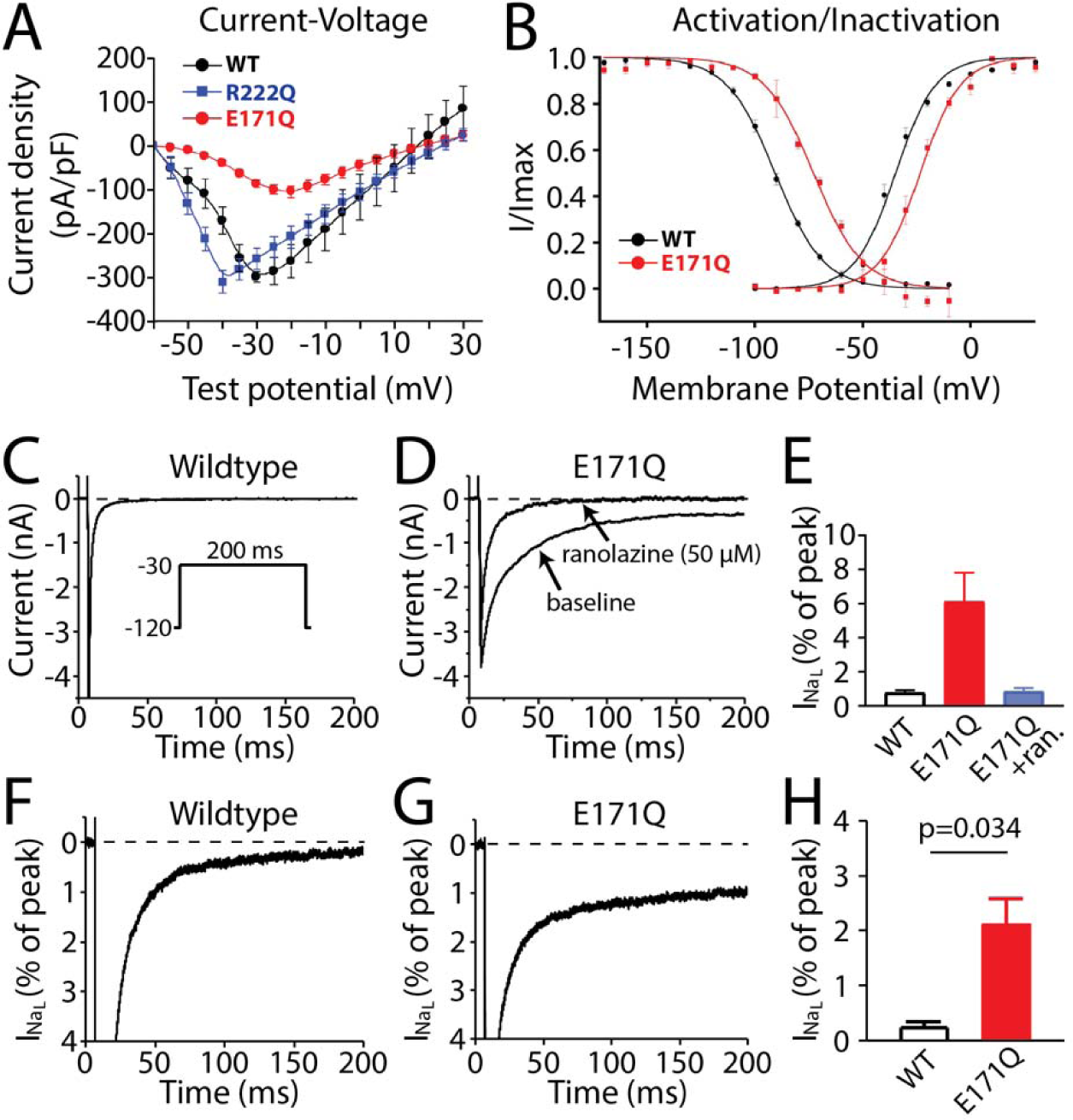
Electrophysiological properties of E171Q in HEK cells and in iPSC-CMs. Panels A-E show HEK293 cells and panels F-H show iPSC-CMs. A) Current-Voltage relationship for WT, R222Q, and E171Q. E171Q has reduced peak current. B) Activation and inactivation curve for WT and E171Q. E171Q has right-shifted activation and inactivation curves. C) Wild-type sodium current (HEK cell). The clamp protocol is shown in the inset. D) E171Q shows enhanced late sodium current which is inhibited by the late current blocker ranolazine. E) HEK cell late current summary data. P<0.0001 overall, with P=0.2 for WT vs E171Q and P=0.09 for E171Q vs E171Q with ranolazine by Dunnett’s test. F-H) Sodium current in population control iPSC-CMs, using the same protocol as in D and E. G) E171Q in an iPSC-CM isogenic to the control. H) Late current summary data in iPSC-CMs.

E171Q reduced peak current magnitude and caused a rightward shift in the voltage dependence of activation and inactivation (Figure 3A-B); the “window” of overlap between the activation and inactivation curves was larger and shifted in a positive direction, in contrast to the negative shift previously reported for R222Q.^2-4,24^ At a depolarizing potential of -30 mV, an enhanced “late” sodium current was observed, and abolished with the late current blocker ranolazine (Figure 3C-E); this result was also seen in E171Q iPSC-CMs (Figure 3F-H).

With pulses to more positive potentials, both variants also generated a time-independent outward current that was not seen in wildtype channels (Figure 4A-C). Replacement of external sodium with N-methyl-d-glucamine (NMDG), a cation that does not enter the central pore, resulted in the loss of the inward current but the maintenance of the outward current in R222Q and E171Q (Figure 4D-F). Figure 4G shows current recorded 200 msec after a depolarizing pulse from -120mV: there was enhanced inward “late” current maximal at -50mV, and a gating pore current at potentials greater than -20mV. Gating pore current magnitudes measured in the presence of NMDG were similar in R222Q and E171Q experiments (Figure 4H). The enhanced late current, the outward (gating pore) current in HEK293 cells, and the positive shift in activation were observed at both the VUMC and SFU sites.

**Figure 4:**
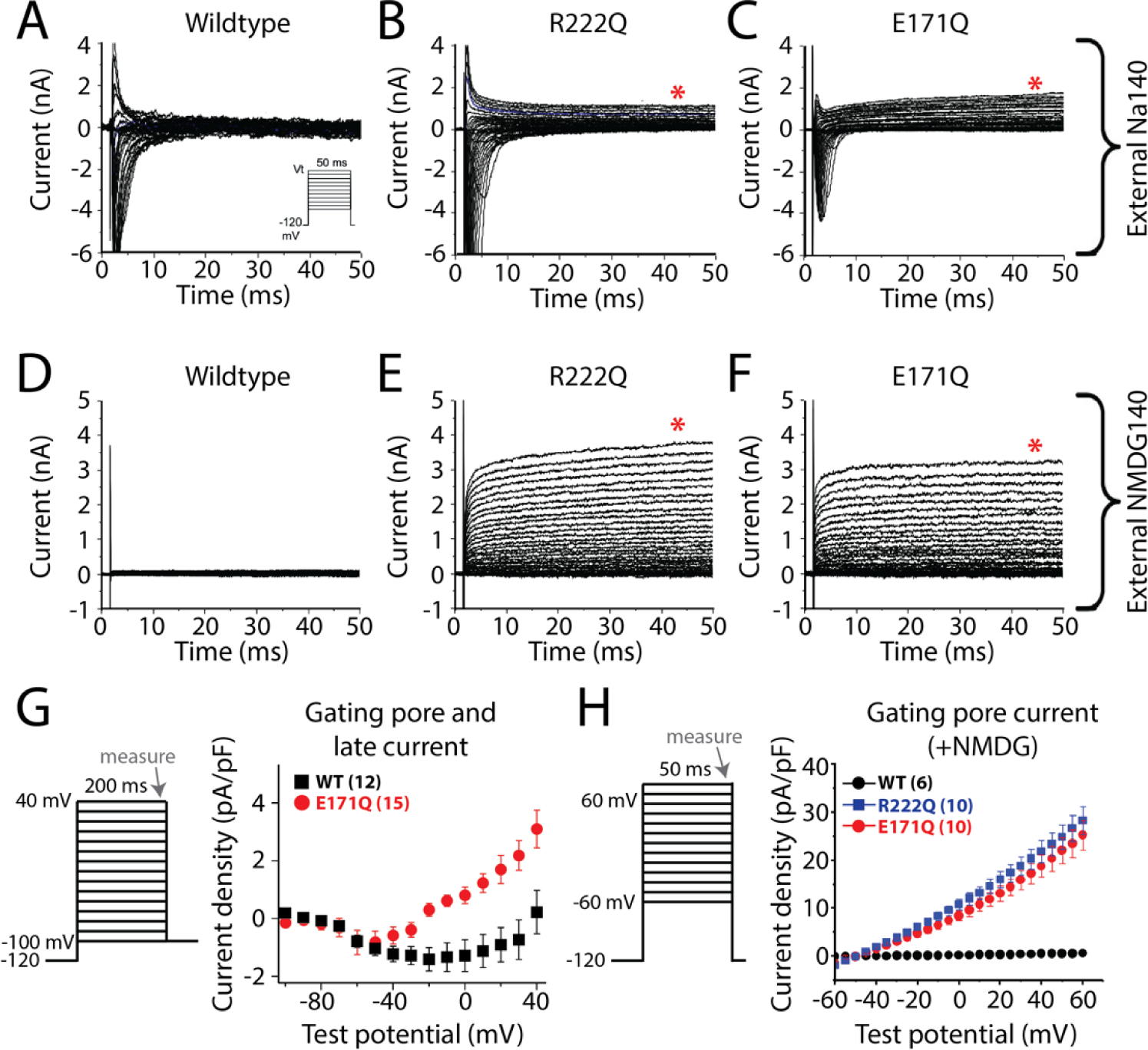
E171Q generates gating pore current in HEK293 cells. A-C) Sodium current traces from *SCN5A* wildtype (A), R222Q (B), and E171Q (C) in an external 140 mM sodium solution. The voltage protocol for A-F is shown in the inset of panel A. Outward pore gating currents are indicated with a red asterisk. D-F) Example sodium current traces from *SCN5A* wildtype (D), R222Q (E), and E171Q (F) in an external 140 mM N-methyl-d-glucamine (NMDG) solution to eliminate inward central pore currents. G) Current density-voltage curve. The data that is plotted is measured at the end of a 200 ms pulse to voltages ranging from -100 mV to 40 mV. H) Gating pore current (quantification of internal NMDG 140 mM traces shown in D-F). The number of studied cells for each mutant in panels G-H is shown in parentheses. G-H) Error bars indicate standard error of the mean.

### Flecainide reduces peak and gating pore currents

In HEK293 cells stably expressing E171Q, 3 μM flecainide abolished both peak current and gating pore current (Figure 5). To evaluate flecainide sensitivity of the gating pore current in the absence of overlapping current through the canonical pore, we used an NMDG-containing extracellular solution (as in Figure 4D-F); under this condition, the IC_50_ for block of the gating pore current was 0.71±0.07 μM (Figure 5G). As shown in Supplementary Figure 1, 1 μM flecainide produced minimal block of peak sodium current.

**Figure 5:**
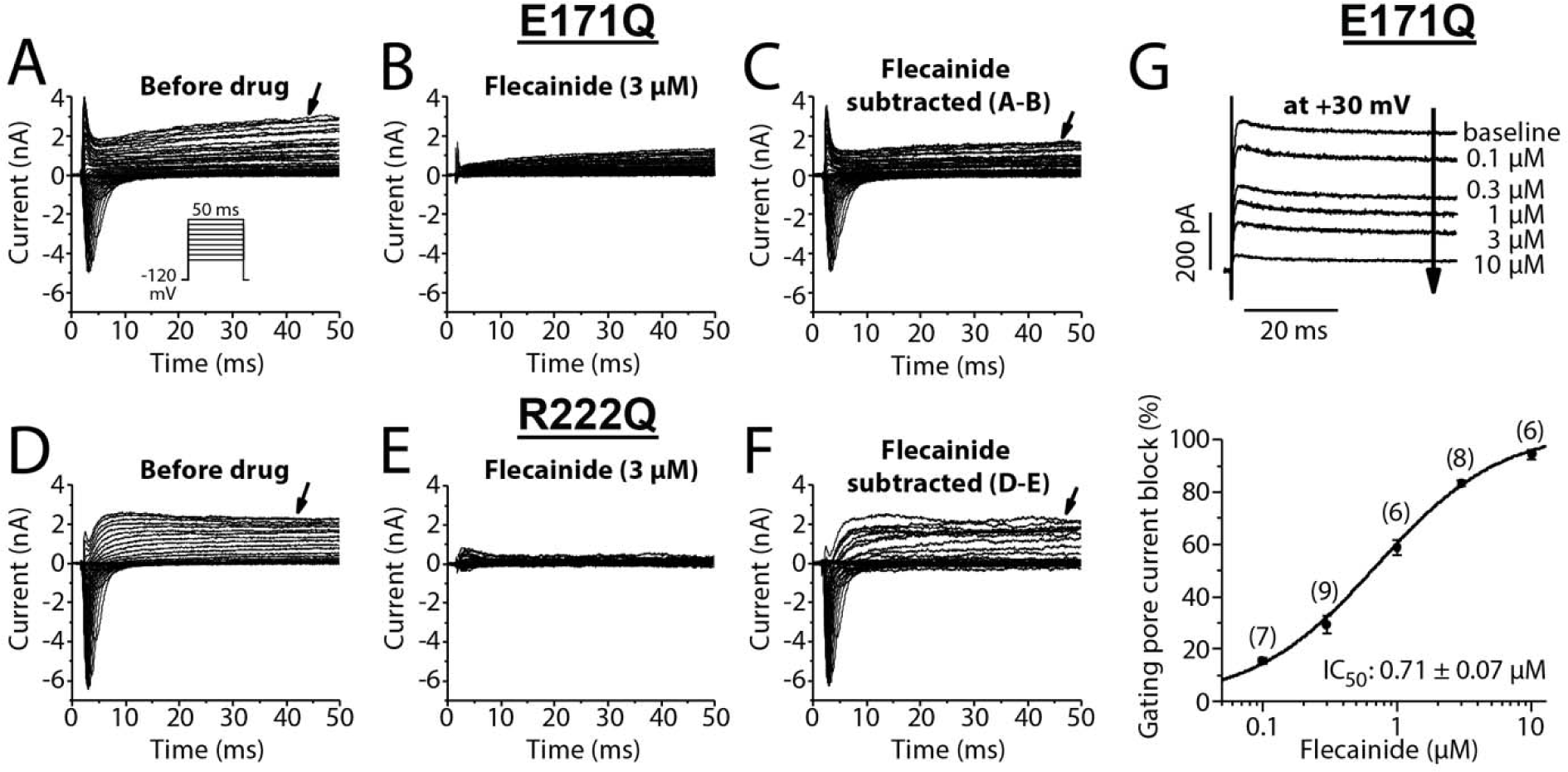
Flecainide effects. A-F) Representative current traces prior to and during flecainide (3 μM) in the two SCN5A variants E171Q and R222Q. Flecainide-sensitive currents were obtained by digital subtraction of current traces prior to and post drug. Panel G is the concentration-response curve for flecainide block of the pore gating current (in the presence of NMDG to block permeation through the canonical pore) in E171Q cells. The inset shows increasing block of gating pore current with increasing concentrations of flecainide. The numbers of cells at the different test concentrations are indicated. Figure S1 shows that 1 μM flecainide had a minimal effect on peak sodium current.

### Gating pore current in E171Q iPSC cardiomyocytes

As in the HEK293 cell experiments, E171Q heterozygous iPSC-CMs also demonstrated outward gating pore currents, whereas the isogenic control cells did not (Figure 6A-C). In iPSC-CMs, action potential durations (APD_90_) in E171Q were significantly prolonged to 779±91 ms (n=12, p=0.0048), compared to 455±30 ms (n=8) in WT cells (Figure 5D, E). In 9/12 E171Q cardiomyocytes, there was beat-to-beat variability in spontaneous depolarization rate, and 3/12 cells showed early afterdepolarizations (Figure 6F).

**Figure 6:**
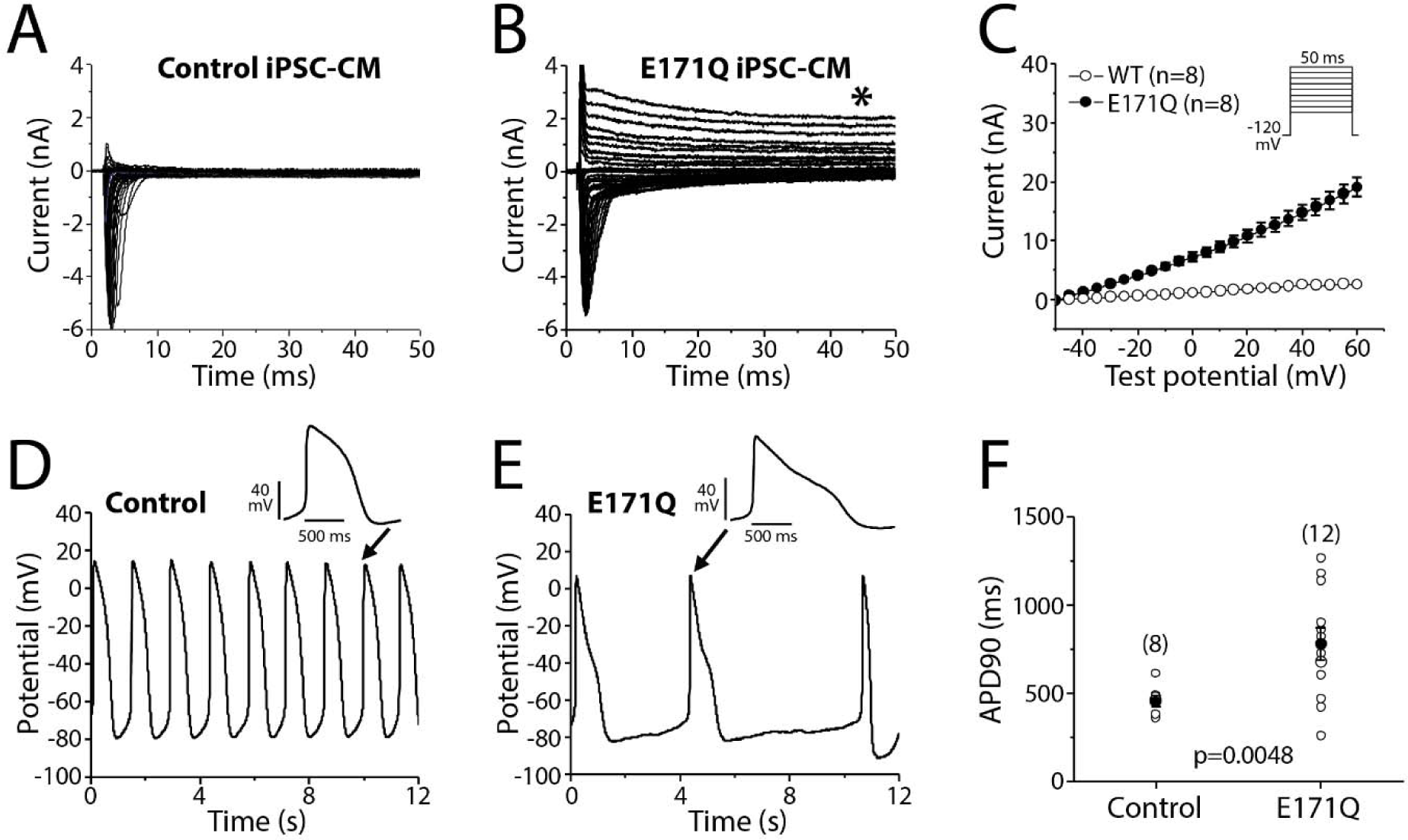
Gating pore current in a CRISPR iPS-cardiomyocyte model of E171Q. A-B) Examples of sodium and pore gating current traces from control (A) and E171Q (B) iPSC-CMs with an external 50 mM sodium solution. The time-independent pore gating current is indicated by a red star (Panel B). Summarized I-V relations for the pore gating currents. The voltage protocol for A-B is shown in the inset of Panel C. D-F) Examples of non-paced spontaneous (D/E) and paced (F) action potentials recorded from two groups of cardiomyocytes. E171Q iPSC-CMs display abnormal (E) and prolonged (F) action potentials. Panel F shows data of APD_90_. The number of studied cells for each group is indicated. Error bars indicate standard error of the mean.

### Molecular dynamics simulations indicate formation of gating pores in E171Q and R222Q but not wild-type channels

We performed μs-scale molecular dynamics simulations of VSD1 (S1-S4) for wildtype, R222Q, and E171Q (n=3 simulations/variant; 1 μs per simulation). Cross sections of simulated proteins showed extracellular and intracellular water-filled clefts that were not connected in the wild-type channel. By contrast, with the R222Q and E171Q variants, the clefts were connected to create a water-filled channel (Figure 7A-C). We took snapshots of the simulations at 1 ns intervals (3000 snapshots in total) and calculated the distances in the simulations between salt bridges involving the mutated residues (E161-R222 and E171-K228; Figure 7D). Wildtype channels spent a large fraction of their time (35.2% of snapshots) with both salt bridges being well maintained, defined here as < 4.0 Å distances between the residue pairs (Figure 7E and Table S3). Overall, wildtype channels spent 93.5% of their time with at least one of the salt bridges being well maintained. By contrast, R222Q resulted in an increased Q222-E161 distance (> 4.0 Å in 99.9% of snapshots) while still maintaining the E171-K228 distance (< 4.0 Å in 90.4% of snapshots) (Figure 7F and Table S4), suggesting that neutralization of the positive charge by variant R222Q disrupted its interaction with E161. Interestingly, E171Q resulted in the disruption of both salt bridges (Figure 7G and Table S5), with 68.8% of snapshots having both distances > 4.0 Å. Thus, the simulations recapitulated the *in vitro* behavior of the channels with respect to generating the water filled channel (Figure 7B, C) that underlies the gating pore current, and suggested that the E171Q variant produced gating pore that was more persistent than that seen with R222Q.

**Figure 7:**
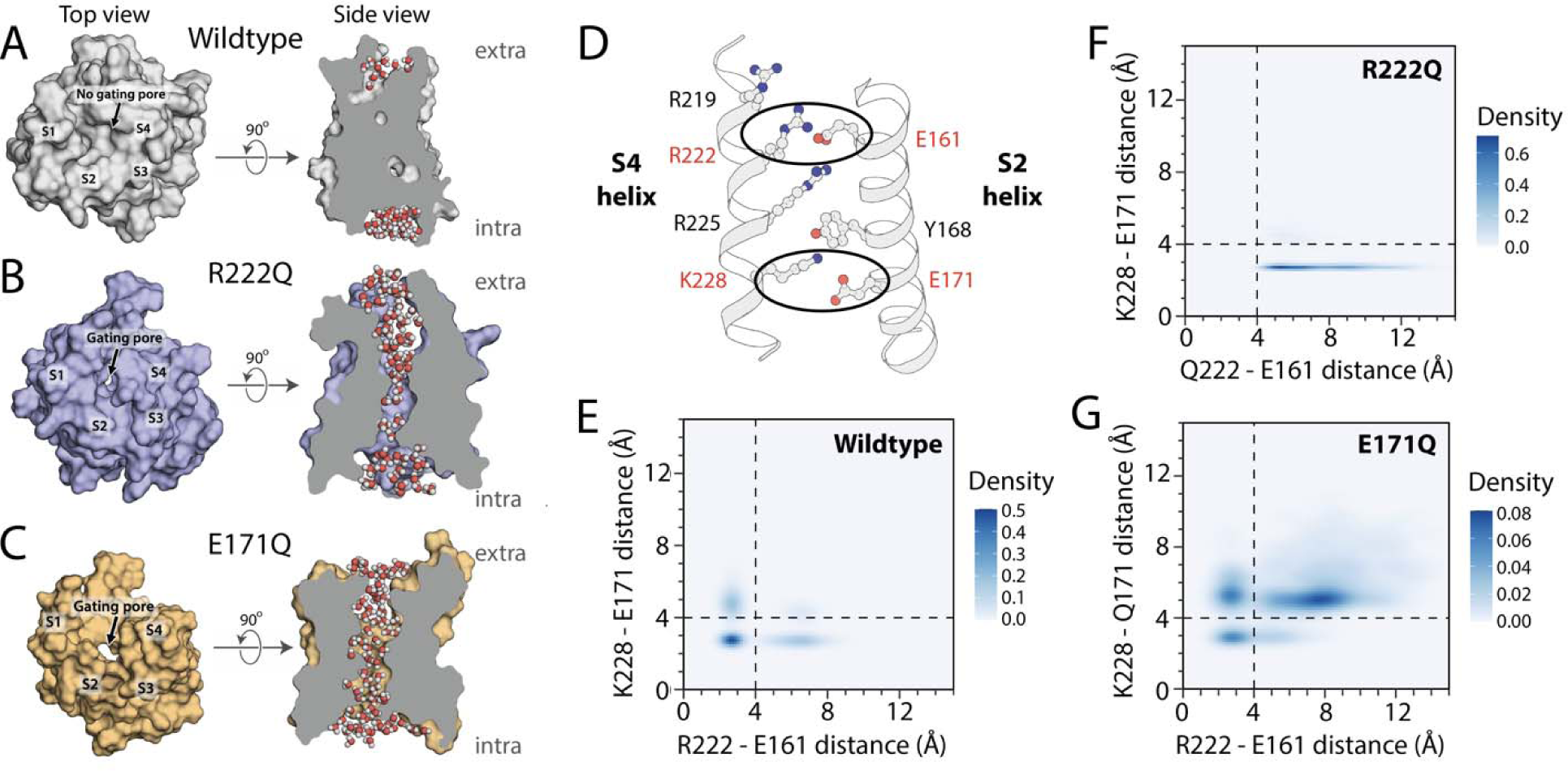
Structural modeling of E171Q and R222Q mutations. A-C) Simulated Domain I voltage sensing domain (S1-S4) of Na_V_1.5. Wildtype (A), R222Q (B), and E171Q (C) structures are shown. Left: view from extracellular region, right: side view. In the extracellular view, wildtype does not have a gating pore, but R222Q and E171Q do. In the side view, water molecules are shown. Both the R222Q and E171Q simulated proteins have water molecules in an extra permeation pathway. D) Ribbon view of the voltage-sensing domain of Na_V_1.5 demonstrating key residues and distances plotted in panels E-G. E-G) Heatmaps of 1) the distance between residues 161 and 222 (x-axis) and 2) the distance between residues 171 and 228 (y-axis). Data are aggregated across all replicate simulations for the entire duration of the simulation, *i.e.* 1000 snapshots captured at 1 ns intervals for each replicate, resulting in 3000 snapshots in total. R222Q specifically increases the 161-222 distance, whereas E171Q increases both distances. See Table S2 for additional simulation details and Tables S3-5 for quantification of the fraction of snapshots in each quadrant in panels E-G.

## Discussion

### Clinical case presentation

We report an infant with frequent ectopy and development of ventricular dysfunction in whom flecainide suppressed arrhythmias and restored normal ejection fraction. While these are the hallmarks of MEPPC, the child also displayed high grade AV block which has occasionally been reported in older subjects^3,4,25^ but is not a common feature of the syndrome. The finding of an *SCN5A* missense mutation was consistent with MEPPC. However, unlike many other mutations reported in this condition, E171Q did not neutralize a positive charge in an S4 helix but rather neutralized a negative charge in S2. At initial presentation, the variant could have been classified as “likely pathogenic” based on AlphaMissense and other *in silico* predictors (PP3, using the ACMG criteria^26-28^), the lack of representation in gnomAD (PM2), and the *de novo* status (PS2). The addition of the functional data we report now results in a classification of E171Q as pathogenic. We cannot rule out the possibility that the *FLNC* and *DMD* VUS may have modified the clinical course, but the correction to normal ejection fraction by flecainide argues that E171Q is sufficient to generate the child’s phenotype.

### Electrophysiological properties of E171Q: comparison to previous studies

The electrophysiologic features of E171Q channels differ somewhat from those reported for R222Q. Cellular electrophysiologic studies of R222Q using heterologous expression reported a consistent negative shift in the overlap between steady state activation and inactivation, generating a larger “window current” at more negative potentials.^2-4^ Similar findings were reported with R225P,^7^ and we also observed this change in cardiomyocytes from R222Q heterozygous mice.^24^ Such a negative shift was predicted to increase sodium influx near the resting potential and thus account for the ectopy characteristic of MEPPC. By contrast, we observe a larger and but positively-shifted window of overlap between steady activation and inactivation curves. The findings thus suggest a common role for the gating pore current, rather than the shift in the window current, in MEPPC. We also find that E171Q generates reduced peak sodium current and this may account for the intermittent AV block.

An increased window current at positive potentials, generating a “late” sodium current, is the hallmark of type 3 long QT syndrome, while QT prolongation and typical polymorphic ventricular tachycardia are not features of the case we report. Moreau et al also observed that the MEPPC-associated mutations R222Q and R225W generated a striking outward gating pore at positive potentials.^10^ This current was originally described in cases of hypokalemic periodic paralysis caused by mutations in voltage sensor arginines in the skeletal muscle channel Na_V_1.4.^29^ Gating pore current in periodic paralysis and MEPPC was most readily detected at positive depolarizing potentials in typical patch clamp experiments but they suggested the pathway could also be open even at negative potentials and could elicit repetitive depolarizations.

R222Q is encoded in *SCN5A* exon 6 but iPSC-CMs express predominantly a fetal splice variant which uses an alternate exon 6; thus usual iPSC-CM culture methods may miss an effect of variants in exon 6.^30^ Recently, Wauchop *et al* used a “biowire” approach to generate mature R222Q iPSC-CMs; they found slightly shorter action potentials than wild-type, and striking contractile dysregulation.^31^ The clinical implications of the contractile abnormality is not clear since suppression of ectopy by flecainide also corrects the ventricular dysfunction in MEPPC. Similarly, Campostrini *et al* have reported that adult type splicing could be restored to iPSC-CMs by culturing them in a three-dimensional microtissue environment and by overexpressing the splicing factor *MBNL1*.^22^

### Proposed structural basis for MEPPC

The Initial mutagenesis experiments in *Shaker* potassium channels that described the GCTC identified a set of negatively charged and hydrophobic residues (including a phenylalanine at the position corresponding to Y168 in NaV1.5; Figure 2) near the voltage sensor that stabilize the mobile S4 helix and prevent a water filled gating pore.^16^ Jiang *et al* proposed a structural model for hypokalemic and normokalemic periodic paralysis in which mutations of arginine residues in sodium or calcium channel voltage S4 sensors generated a water-filled permeation pathway distinct from the ion channel’s pore.^32^ They proposed that mutations in arginines toward to extracellular aspect of S4 generated this gating pore current pathway only in the resting state, but that the pathway was present both in resting and activated states with mutations deeper in the channel.^32,33^ To date, the vast majority of reports of gating pore current as a disease mechanism in MEPPC, and in other ion channel diseases like periodic paralysis, epilepsy, hemiplegic migraine, or malignant hyperthermia, have been confined to charge neutralizing mutations in voltage sensor arginines.^15,34^ One variant, G213D which substitutes a bulky aspartate for a small glycine in the Na_V_1.5 domain 1 S3-S4 linker, has been reported to generate both a large window current and a gating pore current at negative potentials.^13^ G213D iPSC-CMs displayed afterdepolarizations that were corrected with flecainide; whether G213D interacts with the GCTC was not addressed.

The data we present here demonstrates that neutralization of an S2 negative charge that interacts with S4 positive charges can also generate a water filled channel and the MEPPC phenotype. The molecular dynamics simulations suggest that E171 interacts not with an arginine but with the lysine at position 228 in the S4, and that Q171 generates a more persistent and larger gating pore than Q222. Interestingly, the original description of the GCTC in Shaker^16^ suggested that a lysine rather than an arginine at the analogous position near the intracellular face of the channel^15^ plays a critical role in occluding a potential gating pore. The data with extracellular NMDG show that while the peak current (i.e., that through the conventional pore) is abolished, the gating pore current is increased (Figure 4); that finding supports the idea that the E171Q gating pore does not display cation selectivity.

### Limitations

Jiang *et al* identified the flecainide binding site on Na_V_1.5 in the central pore of the channel,^33^ but whether flecainide binding to this site allosterically modifies the gating pore or has a distinct binding site in/near the gating pore region is unknown. The E171Q iPSC-CMs we generated displayed unusually long action potentials and afterdepolarizations. These findings can be explained by the enhanced window current at positive potentials but leave unanswered the question of why QT values in our patient are not unusually long; it may be that the gating pore current serves to limit action potential prolongation in a syncytium.

## Summary

The vast majority of MEPPC cases arise from neutralization of a positive charge in the S4 voltage sensor of the cardiac sodium channel. We report a case of MEPPC due to E171Q, a negative charge neutralization in the S2 helix, that presented with very frequent ectopic beats and decreased ejection fraction, corrected by flecainide. We show that the mutant cardiac sodium channel generates enhanced window current at positive potentials as well as a gating pore current and we use molecular dynamics simulations to demonstrate that E171Q creates a water-filled permeation pathway that underlies the gating pore current. An expanded understanding of the genetic and mechanistic basis for this condition is enabling for effective personalized therapy.

## Acknowledgements

We thank Kara Skorge and Maria Calandranis for assistance growing and genotyping cell lines and Zerubabell Daniel for curating gnomAD allele counts.

## Supplemental Figures and Tables

**Figure S1:**
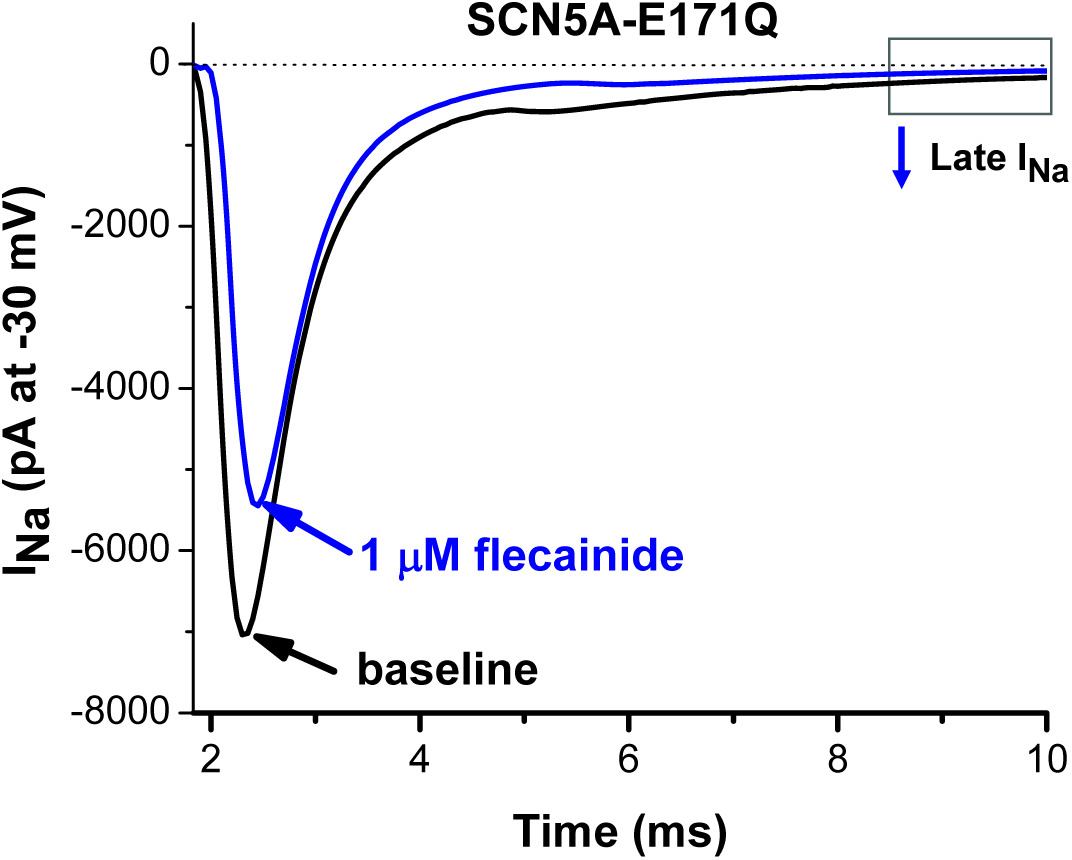
1 μM Flecainide has minimal effect on peak sodium current.

**Table S1:**
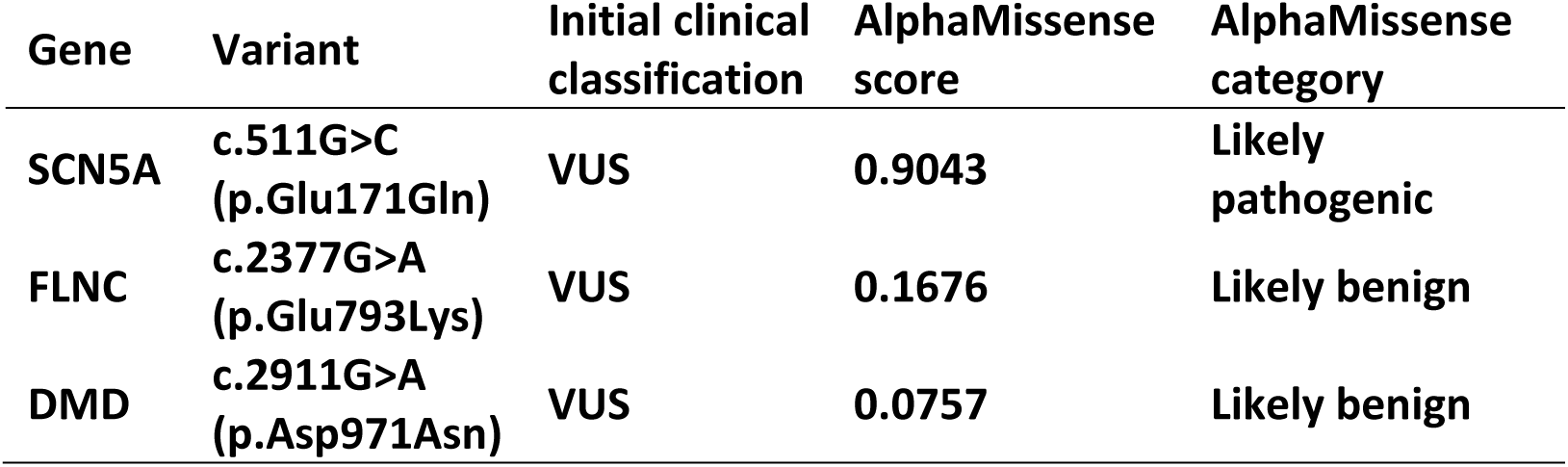
AlphaMissense predictions of variant effects.

**Table S2:** Summary of simulated systems.

| Mutation | Composition | # Atoms | Duration (μs) | # Replicates |
| --- | --- | --- | --- | --- |
| Wildtype | VSD1 (132 residues) + 107 DMPC lipids + water + NaCl | 36,291 | 1.0 | 3 |
| E171Q | VSD1 (132 residues) + 107 DMPC lipids + water + NaCl | 36,381 | 1.0 | 3 |
| R222Q | VSD1 (132 residues) + 107 DMPC lipids + water + NaCl | 36,515 | 1.0 | 3 |

VSD1: Voltage sensor, domain 1
DMPC: Dimyristoylphosphatidylcholine lipid

**Table S3:** Fraction of snapshots for each distance quadrant (Wildtype)

| | E161-R222 ≤ 4.0 Å | E161-R222 $>$ 4.0 Å |
| --- | --- | --- |
| E171-K228 ≤ 4.0 Å | 35.2% | 30.6% |
| E171-K228 $>$ 4.0 Å | 27.7% | 6.5% |

**Table S4:** Fraction of snapshots for each distance quadrant (R222Q)

| | E161-Q222 ≤ 4.0 Å | E161-Q222 $>$ 4.0 Å |
| --- | --- | --- |
| E171-K228 ≤ 4.0 Å | 0.1% | 90.3% |
| E171-K228 $>$ 4.0 Å | 0.0% | 9.6% |

**Table S5:** Fraction of snapshots for each distance quadrant (E171Q)

| | E161-R222 ≤ 4.0 Å | E161-R222 $>$ 4.0 Å |
| --- | --- | --- |
| Q171-K228 ≤ 4.0 Å | 8.5% | 5.2% |
| Q171-K228 $>$ 4.0 Å | 17.5% | 68.8% |

## Supplemental Methods

### Electrophysiologic recordings (VUMC)

To record peak currents, late currents, and gating pore currents in HEK293 cells and iPSC-CMs, two extracellular bath solutions were used. In HEK293 cells, sodium and pore gating currents were recoded in a normal Tyrode’s solution, containing (in mmol/L): NaCl 135, KCl 4, CaCl_2_ 1.8, glucose 10 and HEPES 10, with a pH of 7.4 adjusted by NaOH. The pipette (intracellular) solution contained (in mmol/L) NaF 5, CsCl 120, EGTA 10 and HEPES 10, with a pH of 7.3 adjusted by CsOH. Endogenous outward potassium and chloride currents in HEK-293 cells were eliminated by adding ML-252 (1 μM, to block an endogenous potassium current generated by KCNQ2), 4-aminopyridine (200 μM, to block transient outward K^+^ current I_TO_) and 9-AC (200 μM, to block chloride current).

The pore gating current was also recorded in an external sodium-free and potassium (4 mM)-containing solution that also include NMDG (140 mM). Endogenous currents were blocked as above.

In the iPSC-CM experiments the external Ca^2+^-free solution was modified to lower the sodium concentration to improve voltage control: (in mmol/L) NaCl 50, NMDG 85, glucose 10 and HEPES 10, with a pH of 7.4 adjusted by NaOH. In the experiments to measure the pore gating current only, a Na^+^-free external solution of 140 mM NMDG was used with or /without 4 mM KCl. To record late sodium current in WT- and E171Q-expressed HEK-293 cells, a K^+^- and Ca^2+^-free external solution with 135 mM NaCl was used to optimize late sodium current, which was recorded by 200-ms depolarization pulses to -30 mV from the holding potential of -120 mV. The amplitude of late sodium current was measure at a time window of 195-198 ms before the ending of test pulses. Ten continuous current traces were averaged for analysis of the late sodium current level. A range of concentrations of flecainide (Sigma-Aldrich, USA) was used to test effects on the gating pore currents in HEK-293 cells expressing E171Q or R222Q.

### Action potential recordings (VUMC)

In current-clamp mode, action potentials (APs) in WT- and E171Q-iPSC-CMs were elicited by injection of a brief stimulus current (1-2 nA, 2-6 ms) at a stimulation rate of 0.5 Hz. For AP experiments, the bath (extracellular) solution was normal Tyrode’s, containing (in mmol/L) NaCl 135, KCl 4.0, CaCl_2_ 1.8, MgCl_2_ 1.0, HEPES 5.0 and glucose 10, with a pH of 7.4. The pipette-filling (intracellular) solution contained (in mmol/L) KCl 130, ATP-K_2_ 5.0, MgCl_2_ 1.0, CaCl_2_ 1.0, BAPTA 0.1 and HEPES 5.0, with a pH of 7.3. Ten successive AP traces were elicited with a stimulation rate of 0.5 Hz and averaged for analysis of AP duration at 90% repolarization (APD_90_).

#### Data acquisition (VUMC)

Glass microelectrode recording pipettes with a tip resistance of 0.5 ∼ 1.5 MΩ were used. All data acquisition was carried out using a MultiClamp 700B patch-clamp amplifier and software pCLAMP 9.2. Currents were filtered at 5 kHz (-3 dB, four-pole Bessel filter) and digitized using an analog-to-digital interface (Digidata 1550B, Molecular Diagnostics). To minimize capacitive transients, capacitance was compensated ∼80%. Series resistance was 1-4 MΩ. In all voltage-clamp experiments, the protocols are presented schematically in the figures. Linear leakage currents were digitally subtracted online by the P/N protocol. All ion currents and action potential (AP) parameters were analyzed using pCLAMP9.2 software. All experiments were conducted at room temperature (∼23°C).

### Electrophysiologic recordings (SFU)

Whole-cell patch clamp recordings from Na_V_1.5 expressed in HEK 293T cells were made using an extracellular solution composed of NaCl (140 mM), KCl (4 mM), CaCl2 (2 mM), MgCl2 (1 mM), HEPES (10 mM). The extracellular solution was titrated to pH 7.4 with CsOH. Pipettes were fabricated with a P-1000 puller using borosilicate glass (Sutter Instruments, CA, USA), dipped in dental wax to reduce capacitance, then thermally polished to a resistance of 1.5–2.5 MΩ. Pipettes were filled with intracellular solution, containing: CsF (120 mM), CsCl (20 mM), NaCl (10 mM), HEPES (10 mM) titrated to pH 7.4.^1^

All recordings were made using an EPC-9 patch-clamp amplifier (HEKA Elektronik, Lambrecht, Germany) digitized at 20 kHz via an ITC-16 interface (Instrutech, Great Neck, NY, USA). Voltage clamping and data acquisition were controlled using PatchMaster/FitMaster software (HEKA Elektronik, Lambrecht, Germany) running on an Apple iMac (Cupertino, California). Current was low-pass-filtered at 5 kHz. For peak and late currents, leakage and capacitive currents were automatically subtracted using a P/4 procedure following the test pulse. For measurement of gating pore current with peak current P/4 leak subtraction was not performed. Gigaohm seals were allowed to stabilize in the on-cell configuration for 1 min prior to establishing the whole-cell configuration. Series resistance was less than 5 MΩ for all recordings. Series resistance compensation up to 80% was used when necessary. All data were acquired at least 5 min after attaining the whole-cell configuration, and cells were allowed to incubate 5 min after drug application prior to data collection. Before each protocol, the membrane potential was hyperpolarized to -130 mV to insure complete removal of both fast-inactivation and slow-inactivation.. All experiments were conducted at 22 °C.

### Voltage protocols (SFU)

#### Activation protocols

To determine the voltage-dependence of activation, we measured the peak current amplitude at test pulse voltages ranging from -130 to +80 mV in increments of 10 mV for 19 ms. Channel conductance (G) was calculated from peak I_Na_:

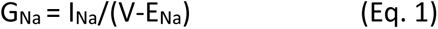

where G_Na_ is conductance, I_Na_ is peak sodium current in response to the command potential V, and E_Na_ is the Nernst equilibrium potential. The midpoint and apparent valence of activation were derived by plotting normalized conductance as a function of test potential. Data were then fitted with a Boltzmann function:

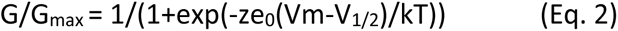

where G/G_max_ is normalized conductance amplitude, Vm is the command potential, z is the apparent valence, e_0_ is the elementary charge, V_1/2_ is the midpoint voltage, k is the Boltzmann constant, and T is temperature in K.

To determine the presence of gating pore current, we performed an offline subtraction of the linear leak, fit using current measured at end of activation epoch between -100 mV and - 60 mV and extrapolated to all voltages.

#### Steady-state fast inactivation protocols

The voltage-dependence of fast-inactivation was measured by preconditioning the channels to a hyperpolarizing potential of -130 mV (to insure complete channel availability) and then eliciting pre-pulse potentials that ranged from -170 to +10 mV in increments of 10 mV for 500 ms, followed by a 10 ms test pulse during which the voltage was stepped to 0 mV. Normalized current amplitude as a function of voltage was fit using the Boltzmann function:

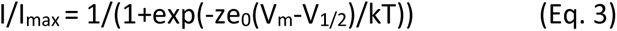

where I_max_ is the maximum test pulse current amplitude. z is apparent valence, e_0_ is the elementary charge, Vm is the prepulse potential, V_1/2_ is the midpoint voltage of SSFI, k is the Boltzmann constant, and T is temperature in K.

#### Persistent current protocols

Late sodium current was measured between 45 and 50 ms during a 50 ms or between 145 and 150 ms during a 200 ms depolarizing pulse to 0 mV from a holding potential of −130 mV (to insure complete channel availability). Fifty pulses were averaged to increase signal to noise ratio.^2,3^

#### Molecular dynamics simulations

We used molecular dynamics simulations to investigate the possibility and extent of spontaneous formation of the gating pore in wild-type and variant Na_V_1.5 Voltage Sensing Domain I (VSD1). We started with a structure of Na_V_1.5 bound to the antiarrhythmic drug quinidine (Protein Databank ID 6LQA; resolution 3.30 Å). We extracted a region (residues 119-250) with domain I pre-S1, S1, S2, S3, S4 and S4-S5 linker sequence. We prepared our simulation systems using the Membrane Builder module^4^ of CHARMM-GUI,^5^ a web-based interface that simplifies building of complex biological simulation systems. Briefly, the VSD1 structure was placed at the center of a rectangular box and the minimum distance between the surface of VSD1 and box boundaries was set to 10 Å. The orientation of Na_V_1.5 with respect to the membrane was obtained from the Orientation of Proteins in Membranes database,^6^ which provides pre-computed protein coordinates with respect to the membrane normal. The system was solvated with water molecules and salted with 0.150 mM NaCl.

We performed all molecular dynamics simulations using the 2020.3 release of the GROMACS simulation package.^7^ We used the all-atom CHARMM36 force field to model protein in conjunction with the CHARMM TIP3P water model.^8^ Before the μs-scale production run, the systems were first relaxed to lower energy through steepest descend energy minimization with a convergence of 1000.0 kJ/mol/nm, a step size of 0.01 nm and a maximum of 5,000 allowed steps. We then equilibrated the systems by following the multi-cycle equilibration scheme as prepared by the Membrane Builder module^4^ of CHARMM-GUI.^5^ We used an integration step of 1 fs in all equilibration cycles to ensure better system stability, but 2 fs in production-run simulations to enable better efficiency. We carried out the μs-scale production-run simulations under the NPT ensemble at 303K and 1 bar with the Nosé-Hoover thermostat^9,10^ and Parrinello-Rahman barostat.^11^ The fast smooth Particle-Mesh Ewald (PME) method^12^ was used for calculating electrostatic interactions with a cut-off value of 1.2 nm, and bonds to hydrogen atoms were constrained using the LINCS algorithm^13^ with default parameters. We performed three replicate simulations each of 1 μs long (totalling 3 μs) for each system (i.e. wild-type, R222Q, and E171Q).

